# Evaluation of Association between *CSN2* Genetic Marker 7:32016718C>A with A1/A2 Milk in Pakistani Buffaloes

**DOI:** 10.1101/2022.08.28.505567

**Authors:** Mehwish Mushtaq, Rashid Saif, Muhammad Hassan Raza, Muhammad Osama Zafar, Zubair Ahmed, Sibghat Ullah, Mateen Ahmad, Asma Karim

## Abstract

Milk is an essential part of the diet that plays an important role in human growth and development due to the presence of different components like calcium, fats and proteins. Among six major proteins of the milk, β-casein is one of them which is being investigated here due to the A1/A2 milk genetic variants and its associated gene of *CSN2* in one of the valued species of *Bubalus bubalis* in Pakistan. Fifty buffalo samples were genotyped using the ARMS-PCR technique. *CSN2* gene locus 32016718C>A is located on Chr.7 with CDS position c.350C>A on its 7^th^ exon as per (GenBank transcript ID: XM_006071124.3). In β-casein, 117^th^ amino acid position where histidine (CAT) is responsible for A1 and proline (CCT) for A2 milk phenotype. A2 milk is considered to be healthy for human health due to the absence of bioactive peptide β-casomorphin-7 (BCM7) in comparison with A1 milk. The current results showed that overall, 100% buffalo population is homozygous wild-type (CC) which means no mutant allele is detected in all studied samples therefore, Pakistani buffalo is exclusively producing A2 milk. Furthermore, this work may also be conducted on different buffalo breeds in Pakistan as well as other species of cattle, goat, sheep and camel to explore their genomic architecture for this valued trait.

## Introduction

Milk is a nutrient rich food item produced by the mammal’s mammary gland which is an emulsion of fat and protein in water, along with dissolved carbohydrates, minerals and vitamins. Milk constitutes 80% and 15% of water and proteins respectively. These proteins have anticarcinogenic, antibacterial and immunomodulatory activities. β-casein is one of the important protein making 75-85% of bovine milk protein [1, 2]. Billions of people all around the world consume milk and dairy products and its role in human nutrition has been increasingly discussed over recent years. Mainly cattle, buffaloes, goats, camels and sheep milk is in human consumption, particularly the buffalo alone produced 127 million tons of milk globally in 2018 [3].

In the 1980s, medical researchers started to study whether some proteins in milk (of β-casein) during digestion causes positive or negative effects on human health. The interest to differentiate between A1/A2 variant, is started in 1990s by researches conducted by the scientists in New Zealand [4], they concluded that A1 β-casein variant is involved in prevalence of various chronic diseases like type 1 diabetes (DM-1), arteriosclerosis, coronary heart disease (CHD) and sudden infant death syndrome [4–6]. However, this hypothesis is not completely accepted and approved by the scientific community.

β-casein gene *(CSN2)* in *Bubalus bubalis* is located on 7^th^ chromosomes [7] which contain nine exons with A1/A2 mutation at exon 7 of genomic position g.32016718C>A with CDS position c.350C>A as per (GenBank transcript ID: XM_006071124.3). Different genetic variants have been reported, among all the variants A1/A2 milk phenotype is considered to be more important due to its role in producing β-casein protein as vital protein in milk with health benefits. β-casein is a phosphoprotein containing 209 amino acids with polymorphism at 117^th^ position. The substitution of nucleotide (CCT>CAT) causes the production of mutant β-casein by changing its proline amino acids to histidine [8]. A1 allele encodes for histidine having weak bonds and is easily broken into bioactive peptide β-casomorphin-7 while in A2 allele there is no mutation hence producing normal milk with health benefits.

In the current study, *CSN2* gene variability is evaluated for A1/A2 milk production in Buffaloes. In Asia, countries like Pakistan, where buffalo milk is more preferred therefore we initiated this research effort to genotype 7:32016718C>A locus using buffalo genome assembly (GCF_019923935.1) to investigate the alternative allele frequencies if prevalent in this valued species.

## Materials and Methods

### Sample collection and DNA extraction

The random buffalo breed was selected to study association of *CSN2* gene locus (7:32016718C>A) with A1/A2 milk production in local population. The blood samples of 50 buffaloes were collected from the jugular vein using K3-EDTA vacutainers and store it at 4^°^C till further use. GDSBio (http://www.gdsbio.com/en/)genomic DNA extraction kit was used to extract DNA from buffalo blood samples.

### Primer designing

Based on *CSN2* gene sequence (XM_006071124.3), ARMS-PCR primers were designed using OligoCalc (http://biotools.nubic.northwestern.edu/OligoCalc.html) software. Total of five primers were designed which were categorized as forward common, reverse normal (N) ARMS, reverse mutant (M) ARMS to amplify wild and mutant type regions of targeted sequence. Moreover, two internal controls (IC) primers were designed to amplify sequence of the targeted region. 3**’**-end of reverse ARMS primers were selected for variant analysis. A secondary mismatch was also introduced at 4^th^ nucleotide position from 3’-end of reverse ARMS primer to increase specificity. The details of primers are displayed in table 1.

**Table 1:**
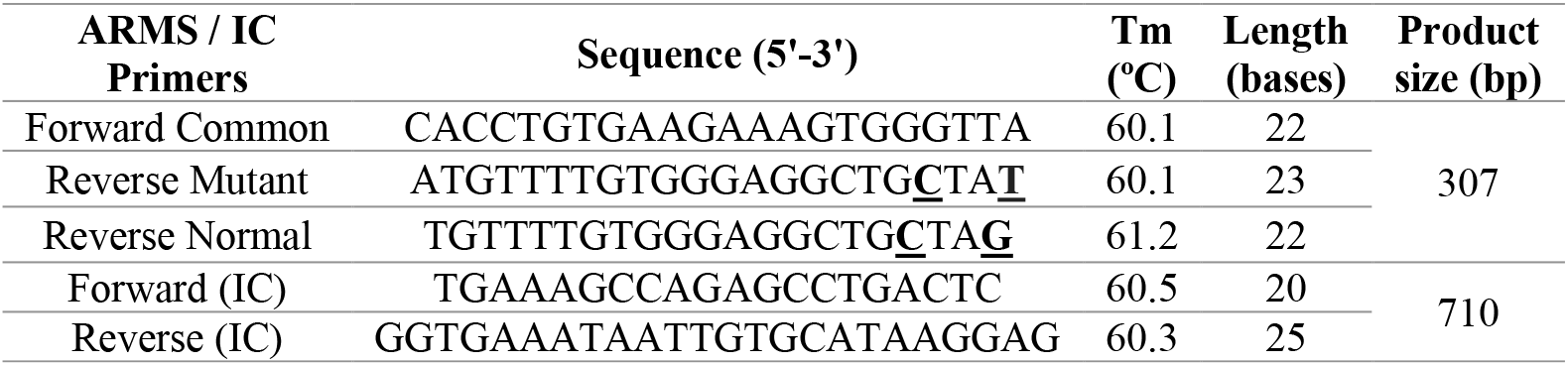
ARMS-PCR primers sequences

### DNA amplification

SimpliAmp thermal cycler was used to amplify wild and mutant type variants. For each sample, two PCR reactions were performed separately with normal (N) and mutant (M) type primers. For normal allele amplification, forward common, reverse N-ARMS and two IC primers were used, while the reaction for mutant allele was carried by using reverse M-ARMS primer.

Reaction mixture of total 16μL was prepared which contains 1μL of extracted genomic DNA, 1μL of 10 mM of each primer (N or M primer, forward common primer, Forward and reverse IC primers), 0.05IU/μL of *Taq* polymerase, 2.5mM MgCl_2_, 2.5mM dNTPs, 1x buffer, and PCR grade water. The PCR protocol was started with initial denaturation at 95^°^C for 5 min, followed by 30 cycles of denaturation (95^°^C for 30s), annealing (60^°^C for 30s) and extension (72^°^C for 30s). After that final extension was at 72^°^C for 10min.

### Statistical association

Normally, Chi-square association test was applied using PLINK data analysis toolset to study association of subject variant with its phenotype.

## Results

### Variant genotyping

A total of 50 buffaloes were screened to analyze the *CSN2* (7:32016718C>A) locus variability for A1/A2 milk phenotype from Pakistan. ARMS-PCR results showed that all the native buffaloes have homozygous wild (A2/A2) genotype which indicates no alternative allele/genotype (A1/A1, A1/A2) in the population. The tendency of homozygous wild-type in *CSN2* gene concluded that all the sampled buffaloes give normal A2 milk. Figure 1 shows the ARMS-PCR amplification gel results of *CSN2* variant along with sampled buffaloes.

**Figure 1:**
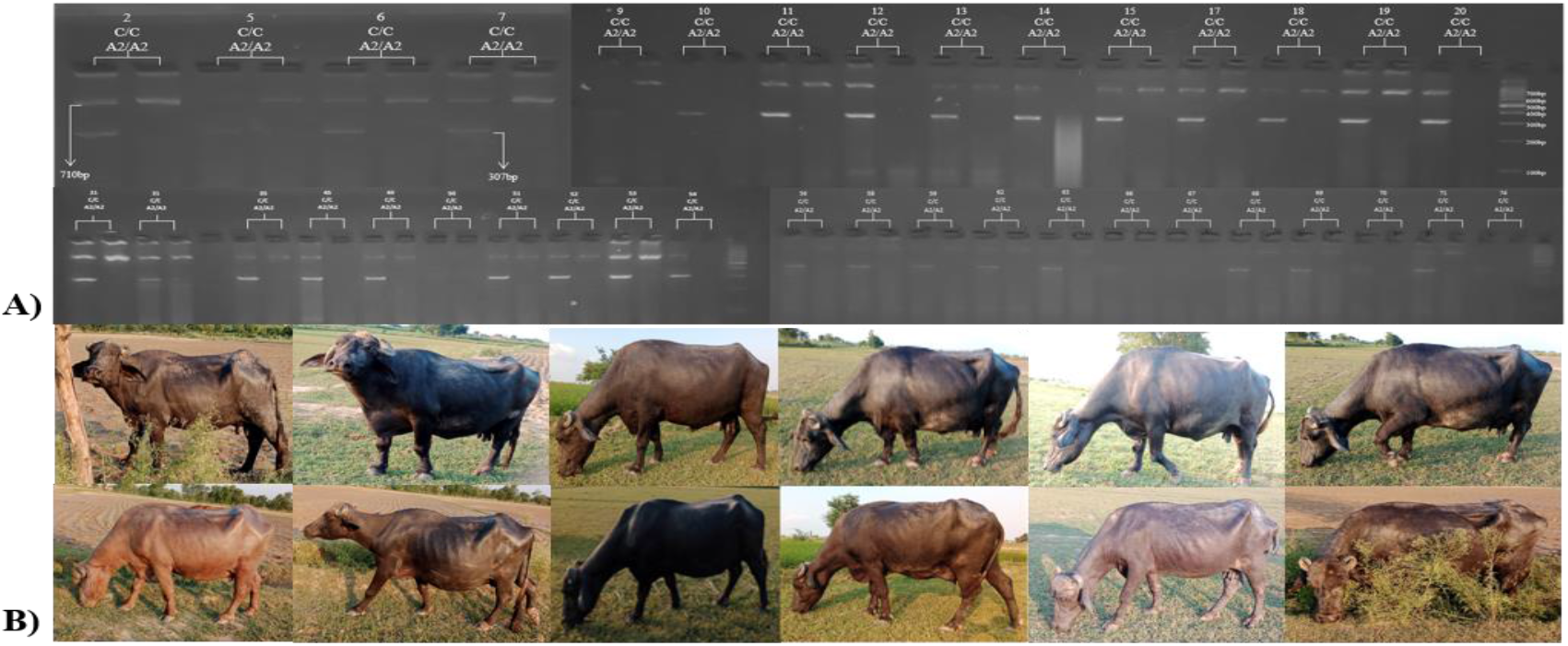
**A)** ARMS-PCR amplification gel results **B)** Sampled buffaloes

### Association analysis

The genotypic percentages, allelic frequencies and chi-square statistical association is calculated using PLINK data analysis toolset. As our 100% of the population is homozygous (CC) with no heterozygous (CA) or homozygous mutant (AA) individuals. Chi-square statistics showed A2/A2 milk (C/C) genotype with allelic frequency of 1, hence Pakistani buffalo population possesses only A2 milk phenotype (Table 2).

**Table 2:** Association analysis of *CSN2* locus in Pakistani buffalo population

| Chr/<br>Position | cDNA variant<br>XM_006071124.3 | Protein variant<br>XP_006071186 | Genotypic Information |  |  | Alternative allele<br>frequency | p-Value |
| --- | --- | --- | --- | --- | --- | --- | --- |
|  |  |  | Homo-wild<br>A2A2/% | Hetero-A1A2/% | Homo- mutant<br>A1A1/% | Native |  |
| NC_059163.1<br>7:32016718 | c.350C>A | p. Pro117His | 1.00/100 | 0/0 | 0/0 | 1.00 | NA |

## Discussion

Cow milk is more preferred in West as compare to Asia, but buffalo milk is mostly consumed especially in South Asia due to its high fat contents despite of its lesser yield. It has more viscosity with composition similar to human milk [5, 9]. Milk proteins including: lactoferricin, glycomacropeptide, casomorphins, lactorphins and casein-phosphopepetides are considered to be vital for human health [2]. The subject variant has been the center of discussion among different researchers as A1 milk causes various health problems. Prevalence of this polymorphism (CCT>CAT) for A1 milk production is more common in cows whereas none of the buffalo was observed with mutant genotype which indicates the fixation of A2 allele in Pakistani buffaloes.

World human population is increasing drastically which is alarming to meet the per capita nutrition requirements. To overcome this problem farming of dairy animals is the best option and is widely adopted all over the world. Several companies were established with the aim of A2 milk production because it is considered healthy than A1 milk. Numerous studies have been conducted for investigating A1/A2 variant in past and further work may be performed to consider the role of other genes and their variants.

The current study has demonstrated the genetic association of *CSN2* variant for A1/A2 milk production in Pakistani buffaloes. In our subject animals, 100% of the buffaloes are homozygous (CC) wild-type having A2/A2 milk phenotype. Similar study was conducted on 66 buffaloes in Ukraine which also showed homozygous A2/A2 genotype in all their screened animals [5]. Another study was conducted in Brazil on 657 buffaloes and all were found with A2/A2 genotype [10]. As for as regional countries are concerned, India has also screened 72 Murrah and Surti buffaloes and revealed the same results of A2/A2 genotypes in their native animals [11].

## Conclusion

*CSN2* gene locus (7:32016718C>A) found with no variability in our sampled buffaloes and displayed 100% homozygous A2A2 genotype producing A2 milk. The current research will be of interest for the researcher, policy maker and ultimately farmers to know the genetic potential of our indigenous buffalos and caveat emptor to introduce exotic animals for enhancing the milk production of our black gold.

## Ethical statement

The ethical statement and approval were exempted for this study. No specific authorization from an animal ethics committee was required. Sampled buffaloes belonged to private buffalo farmers, who joined the present study on a voluntary basis.

## Acknowledgement

The authors acknowledged the buffalo farmers and owners for providing the blood samples.

